# ER discontinuities are common in *C. elegans* neurons, revealing a genetically tractable model for ER network maintenance

**DOI:** 10.64898/2026.03.31.715740

**Authors:** Kelsey N. Mabry, Eric K.F. Donahue, Aaron D. Orgel, Brennen Keuchel, Max G. Kushner, Kristopher Burkewitz

**Affiliations:** Department of Cell and Developmental Biology, Vanderbilt University School of Medicine, Nashville, TN 37240, USA

## Abstract

The neuronal endoplasmic reticulum (ER) extends from the soma into axons and dendrites to coordinate protein trafficking, lipid metabolism, inter-organelle organization, and calcium homeostasis. Conserved genes involved in shaping the tubular ER are implicated in neurodevelopment and neurodegeneration, suggesting that ER structure and dynamics influence neuronal health and drive pathogenesis. However, the links between ER morphology and neuronal function and resilience remain incompletely understood. While models typically depict the neuronal ER as a fully continuous network, here we show that micron-scale ER discontinuities in neurites are unexpectedly common in young, unstressed *C. elegans*. These discontinuities occur in both axonal and dendritic compartments with a consistent frequency that varies between motor and mechanosensory neuron types. Using live imaging and photokinetic assays of endogenously tagged markers of the ER, we confirm that these gaps reflect true loss of ultrastructural continuity. Subpopulations of ER tubule tips are highly motile, and the majority of ER discontinuities are resolved in less than an hour. Suggesting the formation of discontinuities is linked to cellular damage, their frequency increases with both age and environmental stress. Finally, in agreement with prior observations across models, discontinuities are exacerbated by impairment of certain ER shaping factors involved in hereditary spastic paraplegia, such as reticulon. These findings reveal a model where ER discontinuities are not uncommon in healthy animals, and provide a tractable system in *C. elegans* to dissect the molecular mechanisms maintaining ER structural homeostasis in vivo.

## Introduction

The endoplasmic reticulum (ER) is a central hub of protein trafficking, lipid synthesis, membrane homeostasis and calcium signaling. Structurally, the ER forms a dynamic, interconnected network of flattened cisternae and tubules covering virtually all regions of the cell. In neurons, the ER is an especially vast network, extending from the soma into dendritic and axonal subcompartments to support biosynthetic and signaling processes. The extreme anatomy and long distances covered by neurons pose unique challenges for cells to control the spatial organization of the ER, raising questions about how organelle continuity and function is coordinated at long-range.

Classical models generally depict the neuronal ER as a fully continuous network in all cell types, including neurons[1–6]. The soma is enriched with rough ER cisternae and is thought to be the site of the bulk of ER protein synthesis and translocation[4,5]. Outside of the soma, sparser ribosomes are occasionally observed in dendritic branches of some neurons, and these ribosomes are thought to support local translation of certain mRNAs close to their functional sites[7,8]. Predominantly, however, neurites are populated by smooth ER tubules[3–5,9]. While larger axons contain simple ER networks, smaller axons sometimes hold a single ER tubule[4,9]. These tubular subdomains provide local hubs to support lipid and membrane homeostasis, act as a reservoir to buffer and shape Ca^2+^ waves, and engage in contact site-dependent communication with other organelle networks[3–6].

Importantly, the specialized architecture of the neuronal ER is coupled to neuron function and vulnerability. Mutations in conserved genes involved in ER shaping and ER-cytoskeletal coupling are strongly linked to neurodevelopmental and neurodegenerative diseases, most notably hereditary spastic paraplegia (HSP)[10–13]. These factors include the ER fusion factor atlastin, tubule-stabilizing reticulons and receptor expression enhancing protein (REEP)-family members, and the ER-resident, microtubule-severing enzyme spastin[4,12]. Despite these genetic links between ER shaping factors and pathophysiology in the nervous system, however, whether any specific abnormalities in ER structure or organization are sufficient to trigger neurodegeneration remains unresolved. Studies of HSP-associated mutations have shown that their disruption can produce gaps in the axonal ER network, suggesting that loss of ER continuity could be linked to neuronal dysfunction[14,15]. Similarly, prior work showed that atlastin mutations impair the ability of ER tubules to extend into distal dendritic networks, resulting in dendrite branches that are completely unoccupied by the ER[16]. Because ER continuity and extension into distal neurites is thought to be important for long-range diffusion and coordinated calcium handling, these ER gaps and unoccupied neurites in HSP mutants suggest plausible upstream mechanisms of pathogenesis. Intriguingly, EM reconstructions in vertebrate models have revealed occasional gaps in the axonal ER of healthy wild type animals as well[9], though their prevalence remains unclear and difficult to quantify.

Despite the emerging importance of ER morphology in neurons, live-imaging constraints in animal models have challenged systematic visualization and quantitation of ER morphology and continuity directly in the nervous system. Towards this end, invertebrates such as Drosophila and *C. elegans* have been foundational in characterizing the ER in developmental contexts [14–16], but the life-long transparency of *C. elegans* also offers the additional opportunity to examine neuronal ER dynamics through adulthood and aging, when many neurodegenerative conditions manifest. Like vertebrate models, *C. elegans* exhibit clear structural and functional segregation of rough ER into soma and smooth ER tubules into neurite compartments[2], with evidence supporting occasional local translation occurring in neurites[17,18]. Furthermore, EM of *C. elegans* neurons has typically revealed neurites containing just a single ER tubule[2,18], enabling direct and unambiguous observation of ER tubule dynamics which are otherwise difficult to resolve with live-imaging techniques when multiple tubules are present.

Here we use endogenous markers and live-imaging of intact adult animals to characterize ER dynamics in a subset of *C. elegans* sensory and motor neurons. We find that ER discontinuities within primary neurites are surprisingly common under basal conditions, enabling visualization of motile ER tubule tips and resolution kinetics. The prevalence of ER discontinuities increases with age and environmental stress and is further enhanced by conserved ER-shaping factors known to be primary risk factors for HSP and other neurodegenerative conditions. Together these observations establish that neuronal ER continuity is an actively maintained and dynamic property *in vivo* while providing a model for defining how failures in ER structural maintenance arise under physiological conditions.

## Results

### ER membrane and luminal markers reveal discontinuities in *C. elegans* neurites

To visualize ER dynamics in physiological contexts, we previously generated transgenic *C. elegans* with endogenous fluorescent labeling of the ER membrane protein, reticulon/RET-1 [19]. Importantly, this labeling strategy circumvents ER structural and localization artifacts from overexpression, while providing enhanced visualization of ER tubules[19,20]. Through this marker we noticed surprising phenomena in the neurons, including an age-associated increase in the RET-1 levels within neurites[19]. We also occasionally observed what appeared to be potential gaps in RET-1 labeling within neurites (Fig. 1A), which was surprising given the prevailing view that the ER is fully continuous. However, due to the ubiquitous expression of native RET-1 in *C. elegans*, fluorescence signal from the ER of surrounding tissues challenged our ability to visualize and validate these ER discontinuities, necessitating a more targeted approach to labeling and visualizing the ER in neurons.

**Figure 1.**
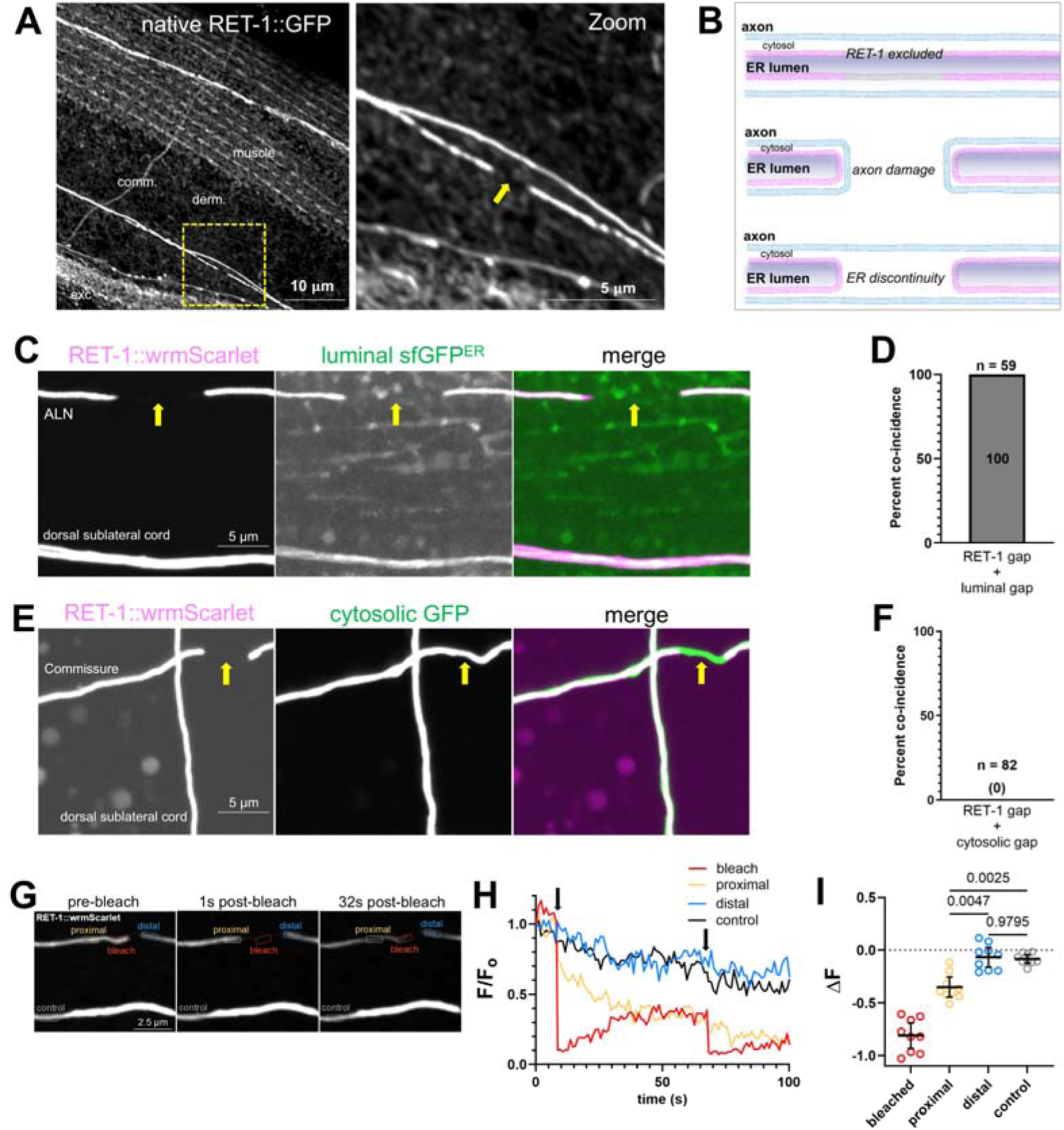
Membrane and luminal markers reveal ER discontinuities in *C. elegans* neurons. A: Maximum-intensity projection from a confocal z-stack of aged (day 7) *C. elegans* expressing endogenously tagged RET-1::GFP. Zoom inset (right) indicates a RET-1::GFP discontinuity in a lateral neuronal process (arrow). Derm. = hypodermis; exc = excretory cell; comm. = motor neuron commissure. B. Potential models explaining gaps in RET-1 labeling. (Top) ER membrane microdomains exclude RET-1: gaps in RET-1 labeling but not in ER luminal markers. (Middle) Axonal damage creates gap: coincident discontinuities in RET-1 and cytosolic fluorescence. (Bottom) Ultrastructural ER discontinuity: coincident gaps in RET-1 and luminal labeling. C. Maximum-intensity projections of ALN and dorsal sublateral cord neurons co-expressing RET-1::wrmScarlet and luminal sfGFP^ER^. D. Percent co-incidence of RET-1::wrmScarlet and luminal sfGFP^ER^ gaps. N = 59 discontinuities. E. Maximum-intensity projections of motor commissure and dorsal sublateral cord neurons co-expressing RET-1::wrmScarlet and cytosolic GFP. F. Percent co-incidence of RET-1::wrmScarlet and cytosolic GFP gaps. N = 82 discontinuities. G. Maximum-intensity projection of RET-1::wrmScarlet fluorescence loss after photobleaching with photobleached (bleach), proximal, distal and control ROIs indicated in frames immediately pre– (left) and post– (right) photobleach. H. Representative trace of RET-1::wrmScarlet intensity over time in each indicated ROI. Arrow marks photobleach. I. Change in mean intensity within each ROI before and after photobleaching. N = 9 neurons from 6 animals, compared by one-way ANOVA with Šídák correction.

To maintain a labeling strategy focused on endogenous ER proteins, we employed a split-fluorescence labeling approach[21,22] to selectively label RET-1 in neurons. Specifically, we knocked wrmScarlet11 into the endogenous *ret-1* C-terminus and drove expression of the complementary, cytosolic wrmScarlet1-10 with the pan-neural *rab-3* promoter. Thus, only the endogenous RET-1 protein in neurons is capable of producing a complete fluorescent wrmScarlet. We next set out to determine whether these apparent gaps represented true discontinuities in the neuronal ER. We reasoned that the likeliest alternative explanations for apparent gaps would include (i) the presence of ER membrane microdomains that exclude RET-1 protein, or (ii) physical damage to the axon’s ultrastructure or anatomy, rather than the axonal ER per se (Fig. 1B). To test the former possibility, we generated animals that co-express neural RET-1::wrmScarlet with superfolder GFP fused with ER signal and retention sequences (sfGFP^ER^), thus serving as an ER luminal marker[23]. Notably, we employed a single-copy knock-in approach to generate this line[24], such that this generic luminal marker complements our endogenous RET-1 labeling strategy while minimizing overexpression. We examined animals for RET-1 neurite gaps in this background, defining them conservatively as regions of the neurite at least 0.5 µm long with no observable ER signal. We found that 100% of RET-1 discontinuities (n = 59) coincide with a luminal discontinuity (Fig. 1C-D). This result indicates that ER discontinuities are not specific to either the membrane or luminal compartment of the ER, nor to a specific fluorescent labeling approach. To reveal any disruption to the neurite ultrastructure, we separately co-expressed pan-neuronal RET-1::wrmScarlet with a cytosolic GFP. However, we found that no RET-1 discontinuities (n = 82) overlapped with gaps in cytosolic GFP (Fig. 1E-F), indicating that RET-1 gaps do not correspond with gross anatomical abnormalities in the neuron itself. Together, these results strongly suggest that the apparent gaps in RET-1 labeling represent true and specific discontinuities in the ER ultrastructure. Though we have previously demonstrated the ability to resolve individual ER tubules *in vivo* in *C. elegans* with confocal or super-resolution microscopy[19], a third possibility could be that fine ER structures in the neurites are too dim to resolve from background fluorescence. To account for this, we performed a fluorescence loss in photobleaching (FLIP) experiment where we photobleached one branch of an apparent discontinuity and measured the effects on fluorescence of the opposite side (Fig. 1G). If these two branches of the ER are physically disconnected and thus unable to share diffusible proteins, photobleaching of one ER branch should have little or no impact on the distal ER tubule. Indeed, we observe no significant loss of RET-1::mScarlet fluorescence on the distal side of ER discontinuities in these experiments, contrasting with the ∼35% decline on the branch proximal to the photobleaching site (Fig. 1H, I). Collectively, these results confirm that the labeling gaps revealed by our RET-1 and sfGFP^ER^ reporters represent true discontinuities in the neurite ER.

### ER discontinuities are surprisingly widespread in vivo and enhanced by stress

Having validated that our reporter system accurately reflects ER discontinuities, we next aimed to estimate the frequency and scope of this phenomenon *in vivo* in *C. elegans*. Importantly, the stereotyped and fully mapped neuronal atlas of the worm enables systematic comparisons between specific neurons and between distinct animals. While the primary processes of many *C. elegans* neurons run in dense bundles and thus can be difficult to resolve, we identified a subset whose processes are spatially separable from neighbors, thus enabling identification of ER discontinuities that are not obscured by overlapping tracts. These neurons included PLM, the commissural motor neurons (AS/DA/DB/DD/VD), the dendritic arbors of the PVD, and ALM/ALN, which run adjacent (Fig. 2A). This subset provides functional and morphological diversity, containing both motor neuron axons and mechanosensory neuronal processes (PLM, ALM, PVD) of varied size and complexity.

**Figure 2.**
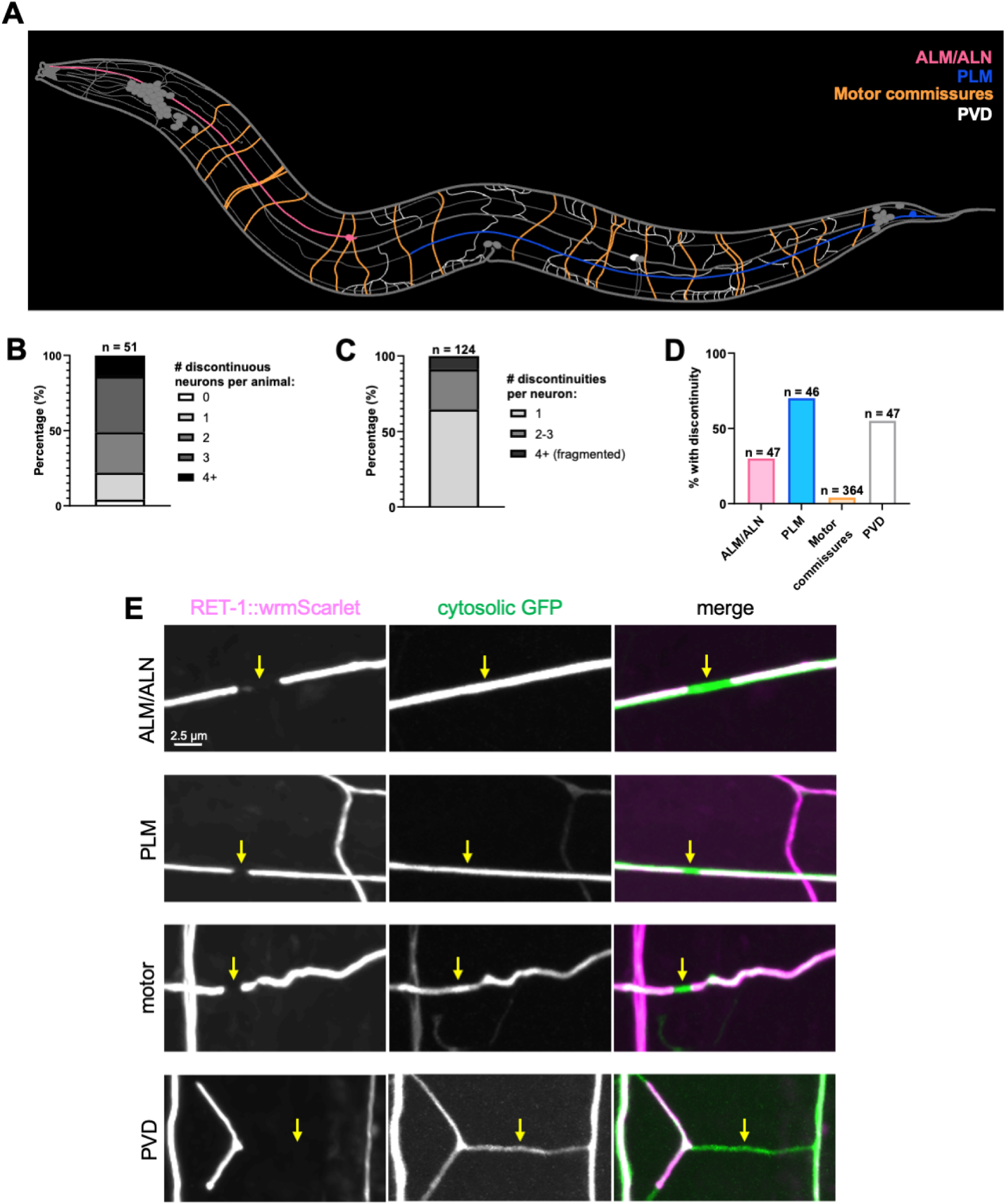
ER discontinuities are common in multiple neuronal subtypes. A: Schematic highlighting the neuronal processes screened for ER discontinuities, including ALM/ALN (pink), PLM (blue), commissural motor neurons (orange), and PVD (white). B: Percentage of animals exhibiting the indicated number of neurons with ER discontinuities. C: Percentage of neurons exhibiting the indicated number of ER discontinuities. D: Proportion of neurons exhibiting at least one discontinuity in healthy, day-1 adults by neuron type. For ALM/ALN, PLM and PVD, n = number of neurons and animals (1 neuron per animal); for motor neurons, n = 364 neurons from 51 animals. E: Max-intensity projections of day 1 ALM (top), PLM (middle top), motor commissure (middle bottom), and PVD (bottom) neurons co-expressing RET-1::wrmScarlet and cytosolic GFP.

To establish a baseline for how common these ER discontinuities are *in vivo*, we manually screened this set of neurons in unstressed, young adult animals. We found that virtually every animal possessed discontinuities in at least one of the screened neurons (Fig. 2B), but few animals exhibited widespread discontinuities across many neurons, indicating these discontinuities occur at a low but consistent rate between animals. This result suggests ER discontinuities are not the result of developmental defects or altered physiological states in certain individuals. We also focused on the neurons that did contain at least one discontinuity to determine the distribution of discontinuities at the cellular scale. We found that ∼60% of those neurons contain a single discontinuity, ∼30% contain 2-3 discontinuities, and ∼10% we categorized as ‘fragmented’ (4+) (Fig. 2C). Thus, discontinuities are most often limited to a single site, but do exhibit a spectrum of severity between individual neurons. The occasional presence of more severely fragmented neurons suggests that there may be rare neuronal states associated with broad-scale ER remodeling or damage. Notably, similar ER fragmentation phenotypes can emerge in mammalian neurons under certain ischemic and depolarizing stimuli[25,26].

Lastly, we assessed how discontinuities were distributed between each of the neuron types examined (Fig. 2D). Interestingly, we found that the sensory neurons, ALM/ALN, PLM and PVD, exhibit discontinuities relatively frequently (∼30-60%), while discontinuities in motor neuron commissures are rarer (<5%). As all these neuronal processes occupy similar subhypodermal tissue environments, differences in mechanical disruption are unlikely to explain the disparity in the frequency of discontinuities. Nevertheless, we sought to test more directly whether physical damage during worm transfer contributes to forming discontinuities. To account for this, we employed gentle eyelash picks to mount the animals, but observed similar results (Fig. S1). Combined with data showing that ER discontinuities are separable from axonal damage (Fig. 1E-F), this result argues against mechanical damage as a primary driver of discontinuities. Thus overall, these findings indicate that ER discontinuities are present in diverse neurons while suggesting that regulation of ER network stability and dynamics varies between neuronal subtypes.

Although we were surprised to find discontinuities at baseline, we also sought to address whether these discontinuities were inducible by environmental and physiological stressors. Previous work in *C. elegans* suggested that heat stress may be sufficient to promote a more fragmented ER network[27]. We tested this systematically by exposing animals to an acute heat stress and quantifying ER morphology in the PLM neuron after a short recovery. Indeed, we observed not just single breaks, but generally a pronounced fragmentation of the ER network in heat-stressed animals (Fig. 3A, B).

**Figure 3.**
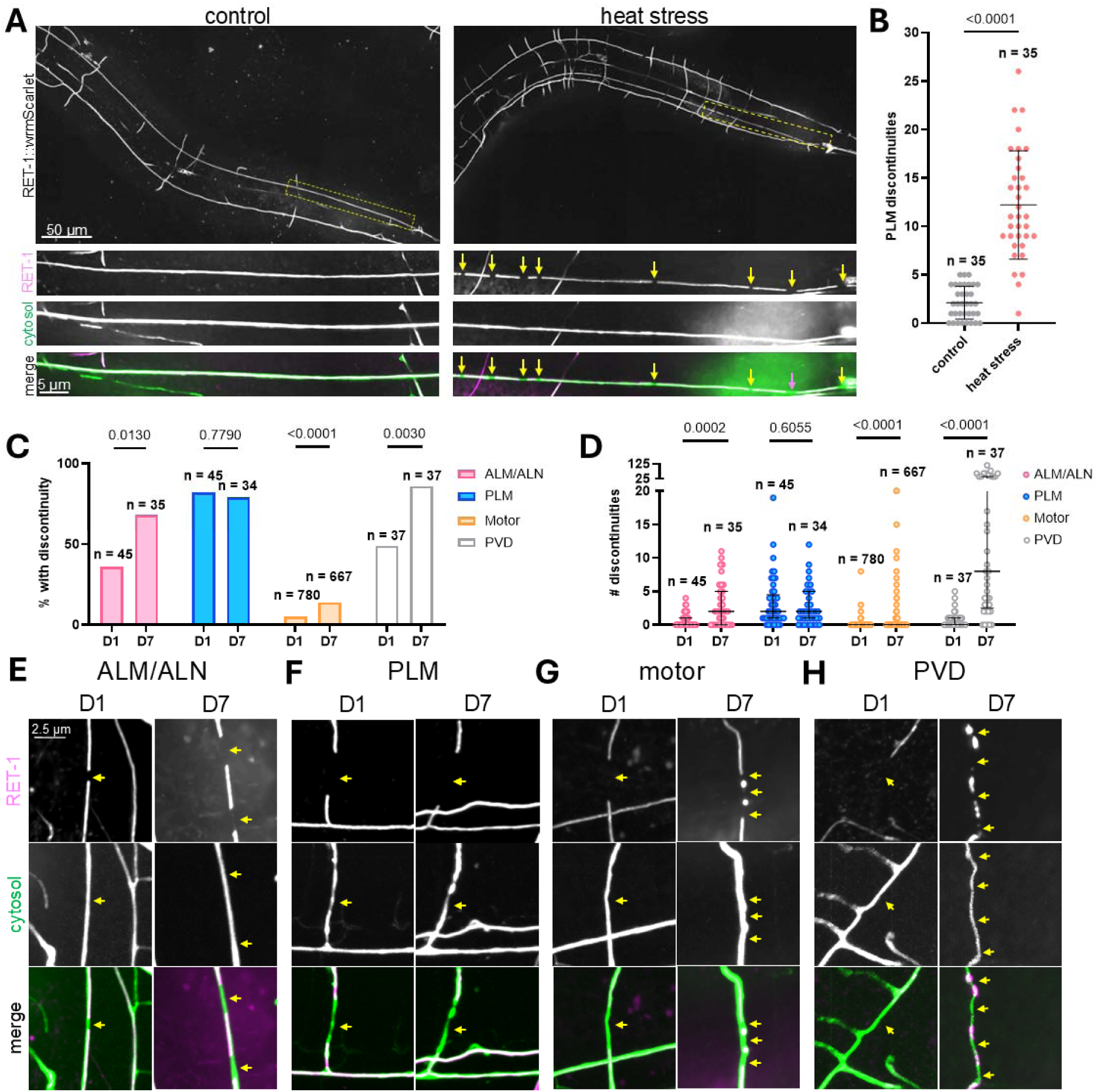
Heat stress and aging enhance ER discontinuities. A: Max-intensity projections of PLM neurons of control (left) and heat-stressed (right) animals co-expressing RET-1::wrmScarlet and cytosolic GFP. B: Number of discontinuities per neuron for control and heat stressed PLM neurons. Data plotted are mean ± standard deviation (SD), compared via Welch’s t-test. C: Proportion of neurons exhibiting at least one discontinuity for day 1 and day 7 neurons by neuron type. Data compared via Fisher’s exact test with Holm-Šídák correction. D: Number of discontinuities per neuron for day 1 and day 7 neurons by neuron type. Data plotted as median ± interquartile range (IQR), compared via Mann-Whitney U-test with Holm-Šídák correction. E-H: Max-intensity projections of day 1 and day 7 ALM (E), PLM (F), motor commissure (G), and PVD (H) neurons co-expressing RET-1::wrmScarlet and cytosolic GFP. For ALM/ALN, PLM, and PVD, depicted n = number of neurons and animals (1 neuron per animal). For motor neurons, n = 780 neurons from 47 animals (day 1) and 667 neurons from 39 animals (day 7).

Aging is also associated with rising ER stress and the onset of neurodegeneration, though how age-dependent changes in ER stress responses and ER morphology are linked remains unclear[28,29]. Leveraging the worm’s unique amenability to live-imaging throughout lifespan, we next asked whether aging alters the prevalence of ER discontinuities. Indeed, we observed a clear increase in the proportion of neurons with discontinuities in aged animals in most neurons examined, including ALM/ALN, motor neurons, and PVD (Fig. 3C). The PLM neurons, which notably exhibit the highest baseline frequency of discontinuities, was the only neuron that did not exhibit an age-associated increase. Similarly, the number of discontinuities present per cell increased across ALM/ALN, motor and PVD neurons in aged animals, but not in the PLM (Fig. 3D). Thus, a subpopulation of neurons with discontinuous and fragmented ER networks increases with age (Fig. 3C-H).

### ER discontinuities reveal tubule dynamics and repair

Because classic models of neuronal cell biology revolve around a fully continuous ER network, we reasoned these discontinuities may be transient in nature. As these discontinuities frequently involve gaps of multiple microns, this model would also predict that free ER tips must be motile in order to reconnect and fuse. To observe ER tip motility, we performed short 5-minute timelapses at 5-7 s intervals. Interestingly, only ∼13% of the discontinuities observed (n = 30) exhibited unambiguous movement during these timelapses, with the remainder of tips appearing to be static. Notably, individual events revealed by live-imaging included clear growth and retraction behavior (Fig. 4A), which we speculate could indicate association with microtubules. We also observed rare events resembling the transfer of a RET-1-labeled particle across the discontinuity (Fig. 4B). Intriguingly, this may be an example of recently identified ER-derived vesicles[30,31]. These distinct vesicles include ribosome-associated vesicles (RAVs), which are ER-derived vesicles thought to act as mobile, translationally competent subdomains of the ER in neurons[30], and REEP-generated vesicles thought to potentially play roles in non-canonical secretion or ER membrane remodeling[31].

**Figure 4.**
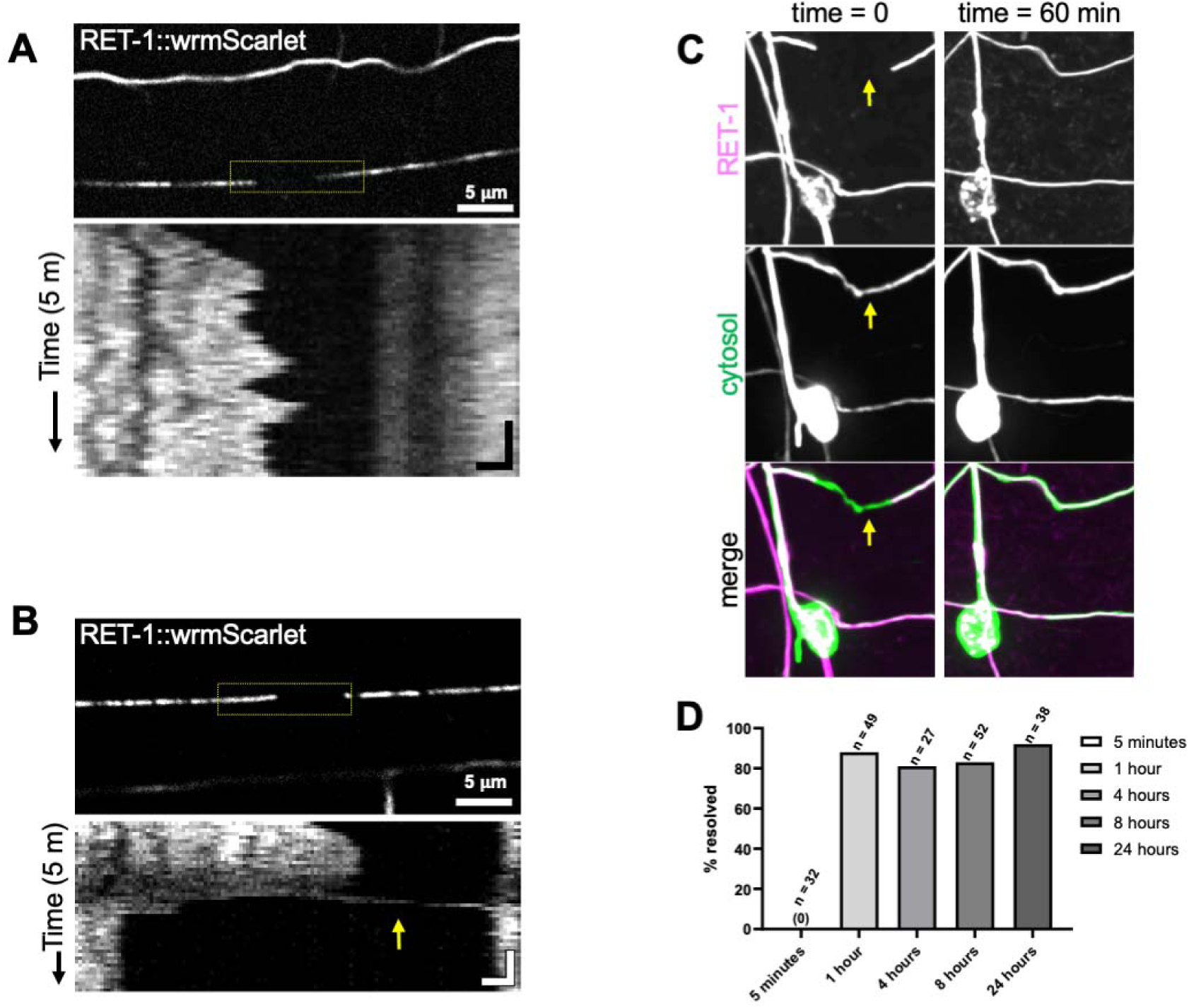
ER discontinuities are dynamic and repaired. A: Timelapse imaging of a PLM neuron harboring a RET-1::wrmScarlet discontinuity with motile tips. Representative image (top) highlights the region used to generate a kymograph (bottom), revealing ER tip growth and retraction (x-axis) over 5 minutes (y-axis). Kymograph scale bars: 1 µm (x), 60 s (y). B: Timelapse imaging of PLM neuron harboring a RET-1::wrmScarlet discontinuity with motile particle and tips. Representative image (top) highlights the region used to generate a kymograph (bottom), which shows ER movement (x-axis) over 5 minutes (y-axis). Arrow marks a RET-1::wrmScarlet particle traveling across the gap. Kymograph scale bars: 1 µm (x), 60 s (y). C: Maximum-intensity projections of ER discontinuity at t = 0 hour (left) and t = 1 hour (right) across RET-1::wrmScarlet (top), cytosolic GFP (middle) and merged channels (bottom). D: Percent of ER discontinuities that resolve by denoted timepoints. N = total ER discontinuities observed for each timepoint.

Next, to determine how long these discontinuities persist, we ran a prolonged timecourse, though with the limitation that we are blind to the precise time at which the gaps we observe were first formed. To minimize phototoxicity, we imaged the discontinuities only twice in this experiment at the start and end points. Consistent with the prediction that these discontinuities would be relatively short-lived, this timecourse revealed that the majority of discontinuities resolve within an hour (Fig. 4C-D). However, across timepoints ranging from 1 to 24 hours, we observed an apparent plateau at ∼80-90% of discontinuities resolving, suggesting a small subpopulation of discontinuities may be persistent (Fig. 4D). Thus, in summary we find that under baseline conditions, only a small fraction of ER tips exhibits extension/retraction behaviors and that ER discontinuities tend to resolve over timescales of less than an hour.

### HSP factors modulate ER discontinuities

Several ER-shaping factors are genetically linked to neurodegenerative disease, of which the best studied example is hereditary spastic paraplegia. However, whether mutations in these ER shaping factors lead to a specific morphological defect in the ER that drives neurodegeneration remains unclear. Prior studies in Drosophila have indicated that loss of hereditary spastic paraplegia-linked factors impact ER morphology in neurons, including causing diffuse ER staining and reduced ER coverage at distal synapses[14,32]. Similarly, atlastin/*atln-1* mutants in *C. elegans* disrupt the ability of ER tips to invade distal dendritic branches in the PVD neuron[16]. More recently, immunostaining of Drosophila axonal ER also revealed seemingly discontinuous, punctate structures in reticulon and REEP mutants[15], similar to what we observe in live *C. elegans* imaging thus far.

We therefore tested whether loss of conserved ER-shaping factors, including reticulon/*ret-1*, atlastin/*atln-1*, REEP5/*yop-1*, and spastin/*spas-1* modulate the formation of discontinuities in *C. elegans.* Because our RET-1::wrmScarlet marker labels the endogenous *ret-1* locus and thus precludes visualization in *ret-1* deletion mutants, we utilized a pan-neuronal, luminal sfGFP^ER^ marker for each of these conditions in the PLM, PVD and motor neurons. In the case of spastin and atlastin/*atln-1* deletion mutants, we observed little consistent or broad effect on the prevalence of ER discontinuities across neuron types (Fig. 5A-D). Surprisingly, however, in PLM neurons atlastin loss resulted in both reduced proportion of neurons affected and the number of discontinuities formed (Fig. 5C, D). Similarly, REEP5/*yop-1* loss resulted in only a mild trend towards increasing discontinuities, reaching statistical significance only in the proportion of motor neurons affected (Fig. 5E, F). In contrast to all other tested HSP mutants, however, loss of *ret-1*, the sole *C. elegans* reticulon gene, resulted in substantial increases in both the prevalence and number of neuronal discontinuities across all tested neuronal subtypes (Fig. 5E, F). Prior work suggested the loss of multiple reticulon and REEP-family members can enhance ER morphology defects[15,33], leading us to test the effect of combined loss of *ret-1* and *yop-1*. While we again observed mild enhancement of discontinuity proportion and number in motor neurons of the double-mutants, the discontinuities appear to be defined overwhelmingly by *ret-1* status (Fig. 5E, F). These results support a model where reticulon/RET-1 plays a conserved and uniquely important role in promoting ER continuity in neurons.

**Figure 5.**
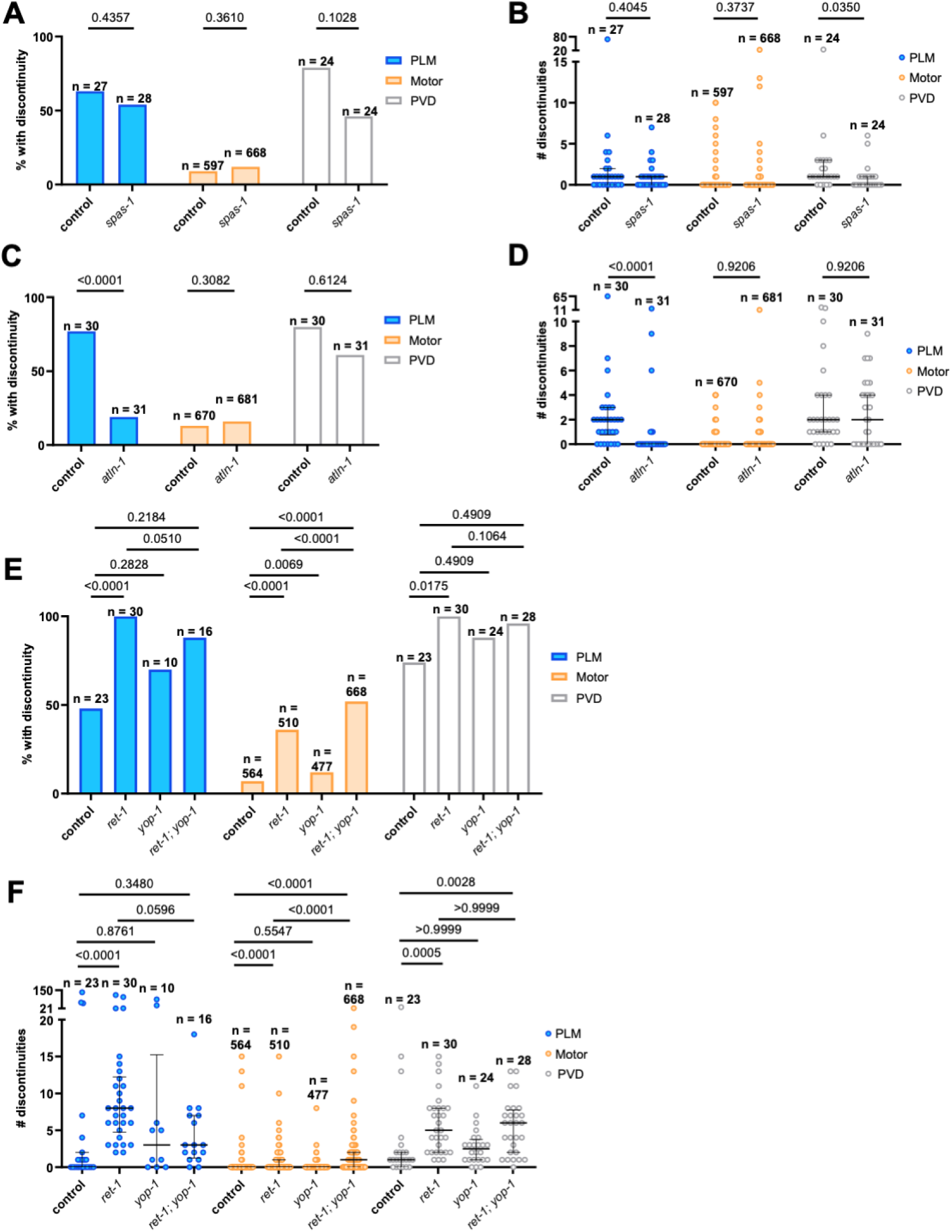
Reticulon mutants exacerbate baseline ER discontinuities. A: Proportion of neurons exhibiting at least one discontinuity for control and *spas-1(ok1608)* by neuron type. B: Discontinuity counts per neuron across control and *spas-1(ok1608)* for indicated neuron types. C: Proportion of neurons exhibiting at least one discontinuity for control and *atln-1(gk5537)* by neuron type. D: Number of discontinuities per neuron across control and *atln-1(gk5537)* by neuron type. E: Proportion of neurons exhibiting at least one discontinuity for control, *ret-1(gk242)*, *yop-1(ok3629)* and *ret-1(gk242); yop-1(ok3629)* by neuron type. F: Number of discontinuities per neuron across control, *ret-1(gk242)*, *yop-1(ok3629)* and *ret-1(gk242); yop-1(ok3629)*. All experiments were performed with the sfGFP^ER^ luminal marker, and data are plotted as median ± IQR in B, D and F. For PLM and PVD the depicted n = number of neurons and animals (1 neuron per animal). For motor neurons, n = 597 neurons from 27 animals (control) and 668 neurons from 28 animals (*spas-1*) in A/B; n = 670 neurons from 30 animals (control) and 681 neurons from 31 animals (*atln-1*) in C/D; n = 564 neurons from 47 animals (control), 510 neurons from 30 animals (*ret-1*), 477 neurons from 27 animals (*yop-1*), and 668 neurons from 28 animals (*ret-1; yop-1*). To compare groups in A, C and E, Fisher’s exact test with Holm-Šídák correction was used; groups in B and D were compared with Mann-Whitney U-test with Holm-Šídák correction. Groups in F were compared via Kruskal-Wallis test with Dunn’s post hoc comparisons.

## Discussion

The neuronal ER is widely modeled as a continuous network, spanning from the soma into dendrites and axons. This continuity underpins our understanding of how neurons maintain homeostasis through long-range transport, calcium signaling and synaptic integrity. Our study leverages endogenous reporters and live-imaging in *C. elegans* to show that it is not uncommon for the ER in neurites to be interrupted by micron-scale gaps, even in healthy young animals. The ER tips at these discontinuities exhibit dynamic behaviors and repair over time, and the prevalence of these discontinuities is dependent upon neuronal subtype, environmental stress and advanced age. Overall, this work highlights a tractable system for investigating the real-time dynamics, regulation and potential pathological consequences of ER network discontinuity across the nervous system.

Our observations of ER discontinuities under baseline physiological conditions lend support to models where ER continuity may be a property that neurons actively control. For example, in hippocampal pyramidal neurons, neuronal activity stimulates ER motility, causing free ER tips to transiently enter and exit dendritic spines[34]. In this model, discontinuities enable free ER tips to explore local neurite anatomy, and activity-dependent enhancement of ER visitation serves as a cell biological mechanism to support synaptic plasticity[6,34]. This exploration may be conserved in *C. elegans,* where ER tubule extension into the complex dendritic arbors of the PVD neuron occurs through a mechanism highly dependent on atlastin function[16]. Similarly, an increase in ER network complexity near neurite branch points can serve as a mechanism to slow cargo and redirect transport[35]. In contrast, however, our study focuses on the discontinuities that occur in the main axonal or dendritic shaft of mature sensory and motor neurons, suggesting a process separable from exploratory ER tip extension at neurite branch points. The simple linear structure of the ER in these regions also raises questions about how these discontinuities form. While there are no established ER fission factors, the loss of continuity in these regions may arise in multiple ways, such as physical associations with microtubules undergoing rapid growth or membrane budding events. Additionally, ER-phagy receptors have recently emerged with key roles in membrane disruption and fragmentation, typically involving the combination of a membrane tether and intrinsically disordered regions[36]. A highly localized accumulation of ER-phagy receptors, which is a role ascribed to multiple reticulon family members[27], could thus potentially trigger localized breaks in the ER tubule as a step in the turnover process. Future studies are needed to delineate molecular mechanisms of ER gap formation and discern whether physiological discontinuities form through processes distinct from stress– or age-induced triggers.

While multiple mechanisms may give rise to discontinuities, their frequency also reflects how efficiently neurons detect and resolve them. Our analysis indicates that the majority of observable discontinuities resolve within an hour, suggesting that maintenance of ER continuity is a constant and efficient process in healthy neurons. However, a small subset of discontinuities seems to persist for extended durations, raising the possibility that certain gaps are refractory to repair or refusion. These unresolved sites may be linked to physical barriers, such as mitochondria or other organelles present within the gap, or lack of access to the machinery required for tip reconnection, such as microtubules or atlastin. Regarding the latter, one of the most surprising results of our study is the finding that null mutants of *atlastin-1* do not exacerbate ER discontinuities in any neuron type examined (Fig. 5). Although the presence of two *atlastin* genes in *C. elegans* creates the potential for genetic compensation, *atln-1* was shown previously to be the dominant regulator of ER morphology in neurons[16,37]. Specifically, hypomorphic or null *atln-1* mutants are sufficient to reduce ER tip invasion into distal PVD dendritic arbors[16] and cause ectopic ER sheet expansion in DA9 motor neurons[37].

Importantly, this result suggests that the mechanisms underlying formation and refusion of ER discontinuities are at least partly distinct from ER tip invasion into dendritic branches. In contrast, *ret-1* mutants exhibited substantial increases in ER discontinuities across all neurons, consistent with prior observations in Drosophila[15]. Because reticulons scaffold and stabilize ER tubule curvature while playing no direct role in membrane fusion, this finding suggests that normal RET-1 levels in the ER membrane act to prevent discontinuities from occurring rather than enhance fusion once gaps have formed. Additionally, the loss of RET-1, one of the most abundant proteins in the smooth ER[38], may induce pleiotropic compensatory changes in the tubular ER that indirectly reduce refusion activities, such as altering cytoskeletal interactions or membrane composition. Additional experiments designed to disentangle the formation and repair processes of ER discontinuities will be required to more precisely define each respective molecular pathway.

Environmental stressors and advanced age are tightly linked with neuronal vulnerability and neurodegeneration, and consistent with these links we also found that heat stress and aging exacerbate ER discontinuities. This finding extends the growing correlation between ER structural features and the pathogenesis of neurodegenerative disease[4,10,11], while providing a powerful model for dissecting the age-associated changes in the neuronal ER that enhance susceptibility to damage and dysfunction. Prior studies have demonstrated clear functional deficits resulting from mutation of HSP genes, including defects in ER calcium buffering, lipid metabolism, microtubule stability, mitochondrial distribution and synaptic function[12–15,32]. Whether and how these functional defects arise from specific structural abnormalities, such as discontinuities, remains unclear, however. The prevalence and amenability of *C. elegans* for finding and observing neurons that contain a discontinuity illuminated in the present study enables more direct association between these gaps and their impact on ER-associated functions. Finally, our finding that multiple HSP-relevant genes in *C. elegans* (spastin, atlastin, and REEP/*yop-1*) failed to enhance discontinuities argues against a simple explanation for these gaps as the driver of HSP pathogenesis. The unifying dysfunction that arises from disruptions in functionally diverse ER shaping factors thus remains murky.

In summary, leveraging the in vivo tractability of *C. elegans* enables real-time observation and dissection of ER dynamics across lifespan and under stress. Our findings advance the classical view of the neuronal ER as an uninterrupted network, revealing instead that neurite continuity is frequently disrupted even under physiological conditions. This new perspective highlights conceptual and practical opportunities to examine how ER integrity is surveilled and maintained across neuronal compartments and to test how ER continuity impacts essential ER functions.

## Methods

### *C. elegans* strains and husbandry

Strains were maintained at 20 °C on 6 cm nematode growth media (NGM) plates seeded with *E. coli* strain OP50-1. Single OP50-1 colonies were cultured overnight in Luria-Bertani (LB) broth at 37 °C and 100 µL of culture was used to seed lawns the following day. Bacterial lawns were grown for 2 days at room temperature before use. Prior to experiments, worm populations were synchronized, and animals were imaged on the first day of adulthood. For experiments involving balanced *atln-1(gk5537)/+* strains, homozygous *atln-1* mutants were isolated as L4s based on *myo-2p*::GFP marker intensity and confirmed to be sterile before imaging experiments as day-1 adults. For aging experiments, animals were transferred every 1-2 days to fresh OP50-1 lawns after reaching adulthood to segregate them from progeny.

The following strains were used in this study: BUZ112 *ret-1(bug25[ret-1::wrmScarlet11]) V; bugEx10[rab-3p::wrmScarlet1-10::unc-54 3’UTR; rab-3p::GFP::unc-54 3’UTR]*, BUZ46 *bugIs9[rab-3p::crt-1(1-17)::sfGFP::KDEL::rab-3 3’UTR] (IV:5015000)*, BUZ279 *bugIs9[rab-3p::crt-1(1-17)::sfGFP::KDEL::rab-3 3’UTR] (IV:5015000); ret-1(bug25[ret-1::wrmScarlet11]) V; bugEx17[rab-3p::wrmScarlet1-10::unc-54 3’UTR]*; BUZ339 *bugIs9[rab-3p::crt-1(1-17)::sfGFP::KDEL::rab-3 3’UTR] (IV:5015000); ret-1(gk242) V*, BUZ340 *yop-1(ok3629) I; bugIs9[rab-3p::crt-1(1-17)::sfGFP::KDEL::rab-3 3’UTR] (IV:5015000)*, BUZ341 *yop-1(ok3629) I; bugIs9[rab-3p::crt-1(1-17)::sfGFP::KDEL::rab-3 3’UTR] (IV:5015000); ret-1(gk242) V*, BUZ344 *bugIs9[rab-3p::crt-1(1-17)::sfGFP::KDEL::rab-3 3’UTR] (IV:5015000); spas-1(ok1608) V*, BUZ346 *atln-1(gk5537[loxP+myo-2p::GFP::unc-54 3’UTR+rps-27p::neoR::unc-54 3’UTR+loxP])/+ IV; bugIs9[rab-3p::crt-1(1-17)::sfGFP::KDEL::rab-3 3’UTR] (IV:5015000)*.

### Cloning and transgenesis

To generate pDB7, the parent plasmid pJA327 (*myo-3p::sfGFP::let-858*) was modified by inserting the N-terminal *crt-1* signal sequence (CRT-1(1-17)) and a C-terminal KDEL ER retention sequence at the ends of sfGFP using fusion PCR. The resulting construct (*myo-3p::CRT-1(1–17)::sfGFP::KDEL::let-858*) was sequence verified. To generate BUZ46 (*rab-3p::sfGFP^ER^*), we employed the SKI-LODGE approach with strain WBM1141 (wbmIs66 [*rab-3p::3xFLAG::dpy-10 crRNA::rab-3 3*′*UTR*] as the genomic insertion background[24]. Repair templates were generated by PCR amplification of the sfGFP^ER^ construct from pDB7 using primers containing 35 bp homology arms corresponding to genomic sequences in the *rab-3* promoter and *rab-3* 3’UTR regions, excluding the rab-3p SKI LODGE 3xFLAG insertion site (5′ arm: AGCCCTATTTTCAGATGACAAGTTTGTACCCCGGG; 3′ arm: AGTTTTTATAGATAGTATAATAGAACGTAGAATTT). The resulting BUZ46 strain was sequence-confirmed. Transgenesis of BUZ112(*ret-1::wrmScarlet11; bugEx10*) was described previously[19].

### Confocal fluorescence microscopy

Synchronized adult animals were manually picked into a ∼2 µL 0.1 µM Polybead suspension (Polysciences) atop a prepared agarose mounting pad (10% agarose in M9 buffer heated and flattened between 2 slides (#1.5, VWR)) and immobilized with a coverslip (#1.5, VWR) as previously described[39]. The pad was further hydrated with M9 applied underneath the coverslip before imaging. Imaging was performed on an Eclipse Ti2 inverted microscope with a Yokogawa CSU-W1 spinning disc (Nikon), a Plan Apo λ objective (20x/0.75 or 100×/1.45) (Nikon), and a Prime 95B sCMOS camera (Teledyne Photometrics). Type LDF immersion oil (Cargille Labs) was used for all experiments. Fluorophores were excited at 488 and 561 nm (dichroic mirror Di01-T405/488/568/647-13×15×0.5 (Semrock)), with respective ET525/36m and ET605/52m emission filters (Chroma). For neural ER projections, imaging was acquired via Z-stack, with capture of the entire neurite width within the image region. For timelapse and fluorescence loss in photobleaching (FLIP) experiments, focal planes were selected at neurite midlines.

### Fluorescence loss in photobleaching

Imaging was performed using a Nikon Eclipse Ti with CSU-X spinning disk scan head (Yokogawa), mini-scanner photomanipulation module (Bruker) and a Prime 95b sCMOS camera (Photometrics). Images were taken with a Apochromat 100x/1.49 NA oil immersion objective at a rate of 1Hz. Bleaching was performed with 5% power of a 100mW 405nm laser (500µs dwell time). Nine images were acquired prior to bleaching a small region at the edge of the observed discontinuity. After bleaching, imaging continued for 60 seconds. The bleaching and imaging process was repeated two additional times. During imaging, care was taken to limit imaging induced stress of the animals. Fluorescence loss in proximal regions and distal regions (across the discontinuity) was measured and normalized to the intensity in the first image in each timecourse. Fluorescence loss in a reference neuron unconnected from the bleached neuron was used to correct for photobleaching caused by imaging. To calculate ΔF for each ROI, the intensity values in the three frames following photobleaching were averaged and subtracted from the mean intensity in the frame immediately preceding the photobleach.

### Screening and quantification of ER discontinuities

Synchronized day-1 adult animals were mounted for confocal microscopy as described above. Leveraging the stereotyped *C. elegans* nervous system, specific identifiable neurons, including the ALM/ALN, PLM, PVD, and commissural motor neurons, were manually screened across the following regions: ALM/ALN process from the midbody ALM soma to the pharynx; PLM process from the lumbar soma to the anterior midbody; commissural motor neurons along their uninterrupted projections between the dorsal and ventral nerve cords; and all PVD secondary, tertiary, and quarternary branches. As *C. elegans* harbors multiple commissural motor neurons asymmetrically distributed on the lateral sides of the animals, the sidedness and the number of motor neurons screened was recorded in each animal. Discontinuities were scored blindly and defined conservatively as unambiguous gaps greater than ∼0.5 µm where the fluorescence intensity was indistinguishable from background. Every scored discontinuity was imaged.

### Heat stress

Synchronized day-1 animals on NGM plates were placed in 37 °C incubators for 1-hour exposure followed by a 1-hour recovery at 20 °C, while control animals were maintained at 20 °C continuously. Animals were mounted and imaged immediately after the 2 hour period.

### ER motility assays

Synchronized day-1 animals were mounted and screened manually for discontinuities in PLM or commissural motor neurons. Upon identification of a discontinuity, a timelapse image series was captured for both GFP and wrmScarlet channels at 5-7 second intervals over a 5 minute period. Resulting series were screened manually for any directed movement during the imaging period and kymographs were generated for select examples with the Fiji (2.16.0/1.54p) Multi-kymograph tool.

### Timecourse of ER re-fusion

Synchronized day-1 animals were mounted and PLM and commissural motor neurons were manually screened for the presence of ER discontinuities. Images were immediately taken of each observed discontinuity to set timepoint 0, and the anatomical location of the discontinuity was precisely recorded. Animals were then rescued from slides to and returned to NGM plates until either the 1, 4, 8, or 24 hour timepoints, when they were re-mounted and imaged again to determine if the discontinuity had resolved. For the 5-minute timecourse, animals were kept on slides. Each discontinuity was only tracked to one timepoint.

### General quantification and statistical analysis

All statistical analyses were performed through GraphPad Prism. To compare proportions of neurons with discontinuities between two conditions, Fisher’s exact tests were applied. To correct for multiple tests across neuron subclasses within each experiment, raw p-values from each comparison were entered into a new table and subjected to Holm-Šidák’s correction with α = 0.05 as the family-wise error rate. Only the adjusted p-values were denoted on graphs. To compare counts of discontinuities, Mann-Whitney U-tests with Holm-Šidák’s correction (2 conditions) or Kruskal-Wallis tests with Dunn’s post-hoc (3+ conditions) were applied. Significance was defined as p < 0.05 based on adjusted p-values.

### Data availability

Strains and plasmids originating in this study are available upon request. Raw image data underlying discontinuity counts is available upon request.

## Acknowledgements

We thank D. Miller, J. Schafer, C. Gailey, R. McWhirter, and S. Kennedy for technical assistance and valuable discussions. To construct the strains used in the current study, N2, WBM1141, VC3086, VC4464, RB1411, and VC441 were provided by the CGC, which is funded by NIH Office of Research Infrastructure Programs (P40 OD010440). Photo-manipulation experiments were performed through the Vanderbilt Cell Imaging Shared Resource, supported by CA68485, DK58404 and EY08126.

## Funding

This work was supported by the Glenn Foundation for Medical Research/American Federation for Aging Research, NIH/NIA R00AG052666 and R01AG073354 (KB), F31AG076290 and NIGMS T32GM007347 (ED).

**Supplemental Figure 1.**
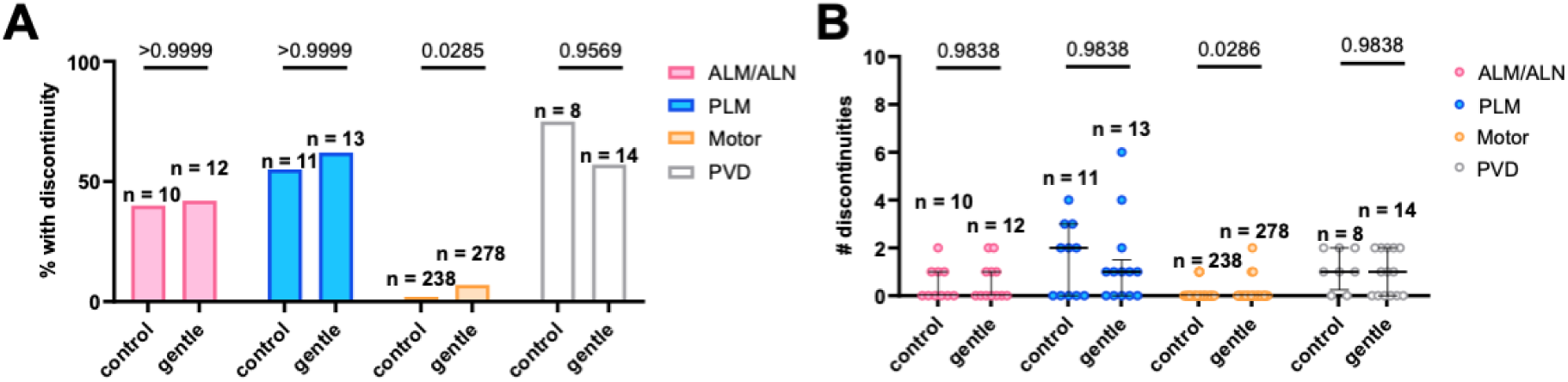
ER discontinuities are not driven by transfer procedures. **A:** Discontinuity proportions for control and gentle-touch ALM/ALN (left), PLM (middle left), motor commissure (middle right), and PVD (right) neurons. Data compared via Fisher’s exact test with Holm-Šídák correction. **B:** Discontinuity counts per neuron for control and gentle-touch ALM/ALN (left), PLM (middle left), motor commissure (middle right), and PVD (right) neurons. Data plotted as exact integers with median ± IQR, compared via Mann-Whitney U-test with Holm-Šídák correction.

## Notes

### Competing Interest Statement

The authors have declared no competing interest.

## References

1. Terasaki M, Slater NT, Fein A, Schmidek A, Reese TS. Continuous network of endoplasmic reticulum in cerebellar Purkinje neurons. Proc Natl Acad Sci U S A. 1994;91: 7510–7514. doi:10.1073/pnas.91.16.7510

2. Rolls MM, Hall DH, Victor M, Stelzer EHK, Rapoport TA. Targeting of rough endoplasmic reticulum membrane proteins and ribosomes in invertebrate neurons. Mol Biol Cell. 2002;13: 1778–1791. doi:10.1091/mbc.01-10-0514

3. Luarte A, Cornejo VH, Bertin F, Gallardo J, Couve A. The axonal endoplasmic reticulum: One organelle-many functions in development, maintenance, and plasticity. Dev Neurobiol. 2018;78: 181–208. doi:10.1002/dneu.22560

4. Sree S, Parkkinen I, Their A, Airavaara M, Jokitalo E. Morphological Heterogeneity of the Endoplasmic Reticulum within Neurons and Its Implications in Neurodegeneration. Cells. 2021;10. doi:10.3390/cells10050970

5. Yperman K, Kuijpers M. Neuronal endoplasmic reticulum architecture and roles in axonal physiology. Mol Cell Neurosci. 2023;125: 103822. doi:10.1016/j.mcn.2023.103822

6. Chanaday NL, Kavalali ET. Role of the endoplasmic reticulum in synaptic transmission. Curr Opin Neurobiol. 2022;73: 102538. doi:10.1016/j.conb.2022.102538

7. Tiedge H, Brosius J. Translational machinery in dendrites of hippocampal neurons in culture. J Neurosci. 1996;16: 7171–7181. doi:10.1523/jneurosci.16-22-07171.1996

8. Kennedy MJ, Hanus C. Architecture and dynamics of the neuronal secretory network. Annu Rev Cell Dev Biol. 2019;35: 543–566. doi:10.1146/annurev-cellbio-100818-125418

9. Terasaki M. Axonal endoplasmic reticulum is very narrow. J Cell Sci. 2018;131. doi:10.1242/jcs.210450

10. Pradhan LK, Das SK. The regulatory role of reticulons in neurodegeneration: Insights underpinning therapeutic potential for neurodegenerative diseases. Cell Mol Neurobiol. 2021;41: 1157–1174. doi:10.1007/s10571-020-00893-4

11. Kulczyńska-Przybik A, Mroczko P, Dulewicz M, Mroczko B. The implication of reticulons (RTNs) in neurodegenerative diseases: From molecular mechanisms to potential diagnostic and therapeutic approaches. Int J Mol Sci. 2021;22. doi:10.3390/ijms22094630

12. Sonda S, Pendin D, Daga A. ER Morphology in the Pathogenesis of Hereditary Spastic Paraplegia. Cells. 2021;10. doi:10.3390/cells10112870

13. Oliva MK, Pérez-Moreno JJ, O’Shaughnessy J, Wardill TJ, O’Kane CJ. Endoplasmic Reticulum Lumenal Indicators in Drosophila Reveal Effects of HSP-Related Mutations on Endoplasmic Reticulum Calcium Dynamics. Front Neurosci. 2020;14: 816. doi:10.3389/fnins.2020.00816

14. O’Sullivan NC, Jahn TR, Reid E, O’Kane CJ. Reticulon-like-1, the Drosophila orthologue of the hereditary spastic paraplegia gene reticulon 2, is required for organization of endoplasmic reticulum and of distal motor axons. Hum Mol Genet. 2012;21: 3356–3365. doi:10.1093/hmg/dds167

15. Yalçın B, Zhao L, Stofanko M, O’Sullivan NC, Kang ZH, Roost A, et al. Modeling of axonal endoplasmic reticulum network by spastic paraplegia proteins. Elife. 2017;6. doi:10.7554/eLife.23882

16. Liu X, Guo X, Niu L, Li X, Sun F, Hu J, et al. Atlastin-1 regulates morphology and function of endoplasmic reticulum in dendrites. Nat Commun. 2019;10: 568. doi:10.1038/s41467-019-08478-6

17. Noma K, Goncharov A, Ellisman MH, Jin Y. Microtubule-dependent ribosome localization in C. elegans neurons. Elife. 2017;6. doi:10.7554/eLife.26376

18. Cuentas-Condori A, Mulcahy B, He S, Palumbos S, Zhen M, Miller DM 3rd. C. elegans neurons have functional dendritic spines. Elife. 2019;8. doi:10.7554/eLife.47918

19. Donahue EKF, Hepowit NL, Ruark EM, Mulligan AG, Keuchel B, Urban ND, et al. ER remodelling is a feature of ageing and depends on ER-phagy. Nat Cell Biol. 2026; 1–16. doi:10.1038/s41556-025-01860-1

20. Shibata Y, Shemesh T, Prinz WA, Palazzo AF, Kozlov MM, Rapoport TA. Mechanisms determining the morphology of the peripheral ER. Cell. 2010;143: 774–788. doi:10.1016/j.cell.2010.11.007

21. Goudeau J, Sharp CS, Paw J, Savy L, Leonetti MD, York AG, et al. Split-wrmScarlet and split-sfGFP: tools for faster, easier fluorescent labeling of endogenous proteins in Caenorhabditis elegans. Genetics. 2021;217: iyab014. doi:10.1093/genetics/iyab014

22. He S, Cuentas-Condori A, Miller DM 3rd. NATF (Native and Tissue-Specific Fluorescence): A Strategy for Bright, Tissue-Specific GFP Labeling of Native Proteins in Caenorhabditis elegans. Genetics. 2019;212: 387–395. doi:10.1534/genetics.119.302063

23. Aronson DE, Costantini LM, Snapp EL. Superfolder GFP is fluorescent in oxidizing environments when targeted via the Sec translocon. Traffic. 2011;12: 543–548. doi:10.1111/j.1600-0854.2011.01168.x

24. Silva-García CG, Lanjuin A, Heintz C, Dutta S, Clark NM, Mair WB. Single-Copy Knock-In Loci for Defined Gene Expression in Caenorhabditis elegans. G3. 2019;9: 2195–2198. doi:10.1534/g3.119.400314

25. Kucharz K, Wieloch T, Toresson H. Potassium-induced structural changes of the endoplasmic reticulum in pyramidal neurons in murine organotypic hippocampal slices. J Neurosci Res. 2011;89: 1150–1159. doi:10.1002/jnr.22646

26. Kucharz K, Lauritzen M. CaMKII-dependent endoplasmic reticulum fission by whisker stimulation and during cortical spreading depolarization. Brain. 2018;141: 1049–1062. doi:10.1093/brain/awy036

27. Serot C, Scarcelli V, Pouget A, Largeau C, Sagot A, El-Hachami K, et al. Reticulon-dependent ER-phagy mediates adaptation to heat stress in C. elegans. Curr Biol. 2025;35: 2365–2378.e7. doi:10.1016/j.cub.2025.04.028

28. Stahon KE, Bastian C, Griffith S, Kidd GJ, Brunet S, Baltan S. Age-Related Changes in Axonal and Mitochondrial Ultrastructure and Function in White Matter. J Neurosci. 2016;36: 9990–10001. doi:10.1523/JNEUROSCI.1316-16.2016

29. Ajoolabady A, Lindholm D, Ren J, Pratico D. ER stress and UPR in Alzheimer’s disease: mechanisms, pathogenesis, treatments. Cell Death Dis. 2022;13: 706. doi:10.1038/s41419-022-05153-5

30. Carter SD, Hampton CM, Langlois R, Melero R, Farino ZJ, Calderon MJ, et al. Ribosome-associated vesicles: A dynamic subcompartment of the endoplasmic reticulum in secretory cells. Sci Adv. 2020;6: eaay9572. doi:10.1126/sciadv.aay9572

31. Shibata Y, Mazur EE, Pan B, Paulo JA, Gygi SP, Chavan S, et al. The membrane curvature-inducing REEP1-4 proteins generate an ER-derived vesicular compartment. Nat Commun. 2024;15: 8655. doi:10.1038/s41467-024-52901-6

32. Summerville JB, Faust JF, Fan E, Pendin D, Daga A, Formella J, et al. The effects of ER morphology on synaptic structure and function in Drosophila melanogaster. J Cell Sci. 2016;129: 1635–1648. doi:10.1242/jcs.184929

33. West M, Zurek N, Hoenger A, Voeltz GK. A 3D analysis of yeast ER structure reveals how ER domains are organized by membrane curvature. J Cell Biol. 2011;193: 333–346. doi:10.1083/jcb.201011039

34. Perez-Alvarez A, Yin S, Schulze C, Hammer JA, Wagner W, Oertner TG. Endoplasmic reticulum visits highly active spines and prevents runaway potentiation of synapses. Nat Commun. 2020;11: 5083. doi:10.1038/s41467-020-18889-5

35. Wang T, Hanus C, Cui T, Helton T, Bourne J, Watson D, et al. Local zones of endoplasmic reticulum complexity confine cargo in neuronal dendrites. Cell. 2012;148: 309–321. doi:10.1016/j.cell.2011.11.056

36. Rudinskiy M, Galli C, Raimondi A, Molinari M. The intrinsically disordered regions of organellophagy receptors are interchangeable and control organelle fragmentation, ER-phagy and mitophagy flux. Nat Cell Biol. 2025. doi:10.1038/s41556-025-01728-4

37. Sun J, Harion R, Naito T, Saheki Y. INPP5K and Atlastin-1 maintain the nonuniform distribution of ER-plasma membrane contacts in neurons. Life Sci Alliance. 2021;4: e202101092. doi:10.26508/lsa.202101092

38. Wang X, Li S, Wang H, Shui W, Hu J. Quantitative proteomics reveal proteins enriched in tubular endoplasmic reticulum of Saccharomyces cerevisiae. Elife. 2017;6. doi:10.7554/eLife.23816

39. Kim E, Sun L, Gabel CV, Fang-Yen C. Long-term imaging of Caenorhabditis elegans using nanoparticle-mediated immobilization. PLoS One. 2013;8: e53419. doi:10.1371/journal.pone.0053419

